# Generative Approaches for Nucleotide Sequences to Enhance Non-coding RNA Classification

**DOI:** 10.1101/2024.11.17.623214

**Authors:** Alisson Gaspar Chiquitto, Liliane Santana Oliveira, Pedro Henrique Bugatti, Priscila Tiemi Maeda Saito, Mark Basham, Roberto Tadeu Raittz, Alexandre Rossi Paschoal

**Affiliations:** Department of Computer Science, Federal University of Technology of Paraná - UTFPR, Corńelio Proćopio, Brazil; Department of Computing, Federal University of São Carlos - UFSCar, São Carlos, Brazil; Laboratory of Bioinformatics, Professional and Technological Education Sector, Federal University of Parańa - UFPR, Curitiba, Brazil; Harwell Science and Innovation Campus, The Rosalind Franklin Institute - RFI, Didcot, United Kingdom

**Keywords:** Data Augmentation, Generative Adversarial Networks, SMOTE, Non-coding RNAs, Artificial Intelligence

## Abstract

Classifying non-coding RNA (ncRNA) sequences, particularly mirtrons, is essential for elucidating gene regulation mechanisms. However, the prevalent class imbalance in ncRNA datasets presents significant challenges, often resulting in overfitting and diminished generalization in machine learning models. In this study, **GEN**erative Approaches for **NU**cleotide **S**equences (GENNUS) is proposed, introducing novel data augmentation strategies using Generative Adversarial Networks (GANs) and Synthetic Minority Over-sampling Technique (SMOTE) to enhance ncRNA classification performance. Our GAN-based methods effectively generate high-quality synthetic data that capture the intricate patterns and diversity of real mirtron sequences, eliminating the need for extensive feature engineering. Through four experiments, it is demonstrated that models trained on a combination of real and GAN-generated data improve classification accuracy compared to traditional SMOTE techniques or only with real data. Our findings reveal that GANs enhance model performance and provide a richer representation of minority classes, thus improving generalization capabilities across various machine learning frameworks. This work highlights the transformative potential of synthetic data generation in addressing data limitations in genomics, offering a pathway for more effective and scalable ncRNA classification methodologies.

## 1 Introduction

The classification of non-coding RNA (ncRNA) sequences, particularly microRNAs (miRNAs) and mirtrons, is essential for understanding gene regulation mechanisms. These small RNA molecules play a critical role in post-transcriptional regulation by modulating messenger RNA (mRNA) levels, thereby influencing a wide range of biological processes [1–5]. The significance of miRNAs research in gene regulation was underscored by the awarding of the 2024 Nobel Prize for their discovery [6, 7].

The successful outcomes presented by the Machine Learning (ML) models have motivated researchers to seek higher accuracy models through complex biological data [8]. However, the classification of ncRNA is often faced by significant class imbalance or small scale in available datasets [3, 9–12]. Both cases leading to overfitting and reduced generalization in machine learning models.

Several strategies have been explored to mitigate this problem, including reducing model complexity, regularization techniques, and acquiring larger training datasets [13–19]. Regularization methods like dropout [13], L1 regularization (lasso) [14], and L2 regularization (ridge) [15] are designed to control model complexity during training. Techniques such as batch normalization and transfer learning have also been employed to accelerate training while preventing overfitting [18, 19]. Additionally, deep learning could better learn from the data but only when the large amount of training set is available. All these approaches often fall short in small-scale applications with limited data, leading to suboptimal generalization.

Data augmentation (DA) offers an alternative by artificially increasing the size of the training dataset, which can improve performance and prevent overfitting. DA has been widely applied in computer vision [8, 20, 21] and natural language processing (NLP) [22–24]. Indeed, applying DA in genome sequence presents unique challenges, in particular to the quality of the generated samples [25]. In image DA, basic transformations such as scaling, rotation, mirroring, or cropping can be applied without altering the original data labels (classes). However, it is not work for biological sequence, particular for RNAs. Given the limited amount of biological data available, strategies to promote the generalization of ML models in genomics are restricted [25].

Generative models (GMs), particularly Generative Adversarial Networks (GANs), have demonstrated remarkable potential in the synthesis of nucleotide and amino acid sequences. For instance, [26] introduced a novel GAN architecture specifically designed to generate synthetic DNA sequences with specific properties. The Feedback GAN (FBGAN) [27] employs a novel feedback-loop architecture that optimizes synthetic gene sequences based on desired characteristics, utilizing an external function analyzer to refine the output. Additionally, [28] developed a deep exploration network aimed at engineering functional DNA sequences. This network effectively explores the input space by minimizing the cost associated with a neural network-based fitness predictor. By integrating a competitive training approach that promotes sequence diversity, they successfully create generators capable of producing thousands of varied sequences. Another notable advancement is ProteinGAN [29], a self-attention-based variant of GANs developed for generating protein sequences with specific functions.

Beyond sequence generation, generative models also play a crucial role in addressing dataset imbalance. For example, [30] explored a GAN-based approach to tackle the issue of data imbalance by leveraging the overall data distribution. In this context, GANs were employed to generate synthetic data that closely mirrors real data, thereby enhancing the performance of machine learning models by mitigating class imbalance in biological sequence analysis. Their study involved four distinct classification tasks utilizing different amino acid sequence datasets.

Among these contributions, [26, 27, 29] focus on applying GANs to generate biological sequences based on nucleotides or amino acids, particularly in creating protein domains. In contrast, [28] concentrates exclusively on the synthesis of DNA sequences. Each of these studies adopts varied methodologies for evaluating the generated samples, with a strong emphasis on assessing the characteristics exhibited by the proteins. Notably, the work of [30] stands out for its commitment to balancing datasets, thereby optimizing classification tasks within the protein domain.

Other approaches that are not based on generative models have been explored in the literature to address class imbalance issues. Among these, Synthetic Minority Over-sampling Technique (SMOTE) is the most used technique nowadays in supervised pre-miRNA classifiers [31]. For example, [32, 33] applied SMOTE for dataset balancing, successfully using it on the resulting matrix from feature extraction. However, there are no studies reporting the application of SMOTE at nucleotide level, for miRNA and mirtron.

This study proposes a novel data augmentation strategy called GENNUS - **GEN**erative Approaches for **NU**cleotide **S**equences. GENNUS harnesses the power of GANs and SMOTE in mirtron and miRNA sequences data analysis. Five approaches are introduced - two based on GANs and three on SMOTE - to address data imbalance in ncRNA classification. The experiments demonstrate that GAN-based methods outperform SMOTE in generating diverse and realistic synthetic ncRNA sequences, significantly improving model performance. Notably, combining real and synthetic data up to a ratio of 2:8 (only 20% real data) enhances classification accuracy. A key advantage of these approaches is their ability to improve model performance without requiring feature engineering, simplifying data preparation. A benchmark was established to compare state-of-the-art mirtron classification tools in addressing the miRNA and mirtron classification problem. This work provides valuable insights into the use of GANs and SMOTE for data augmentation, offering effective solutions to the challenges of data scarcity and imbalance in genomics, particularly for the ncRNA problem.

## 2 Results

Five generative approaches - two GAN-based (WGAN and FBGAN) and three SMOTE-based (SMOTEA, SMOTEB and SMOTEC) - were employed to synthesize data, while seven classification tools were used to evaluate the performance of combinations of real and synthetic data. A series of experiments was conducted, featuring a control group (real data) and experimental groups (real data combined with synthetic data), to evaluate the effectiveness of the proposed approaches. The results of these experiments are presented in the following sections.

### 2.1 Balancing with synthetic data

To investigate whether the synthetic data created by our generative approaches can optimize classification tools, Experiment Design I was employed (Section 4.5.1). Fig. 1A shows the results between control and experimental groups for the seven classification tools.

**Fig. 1.**
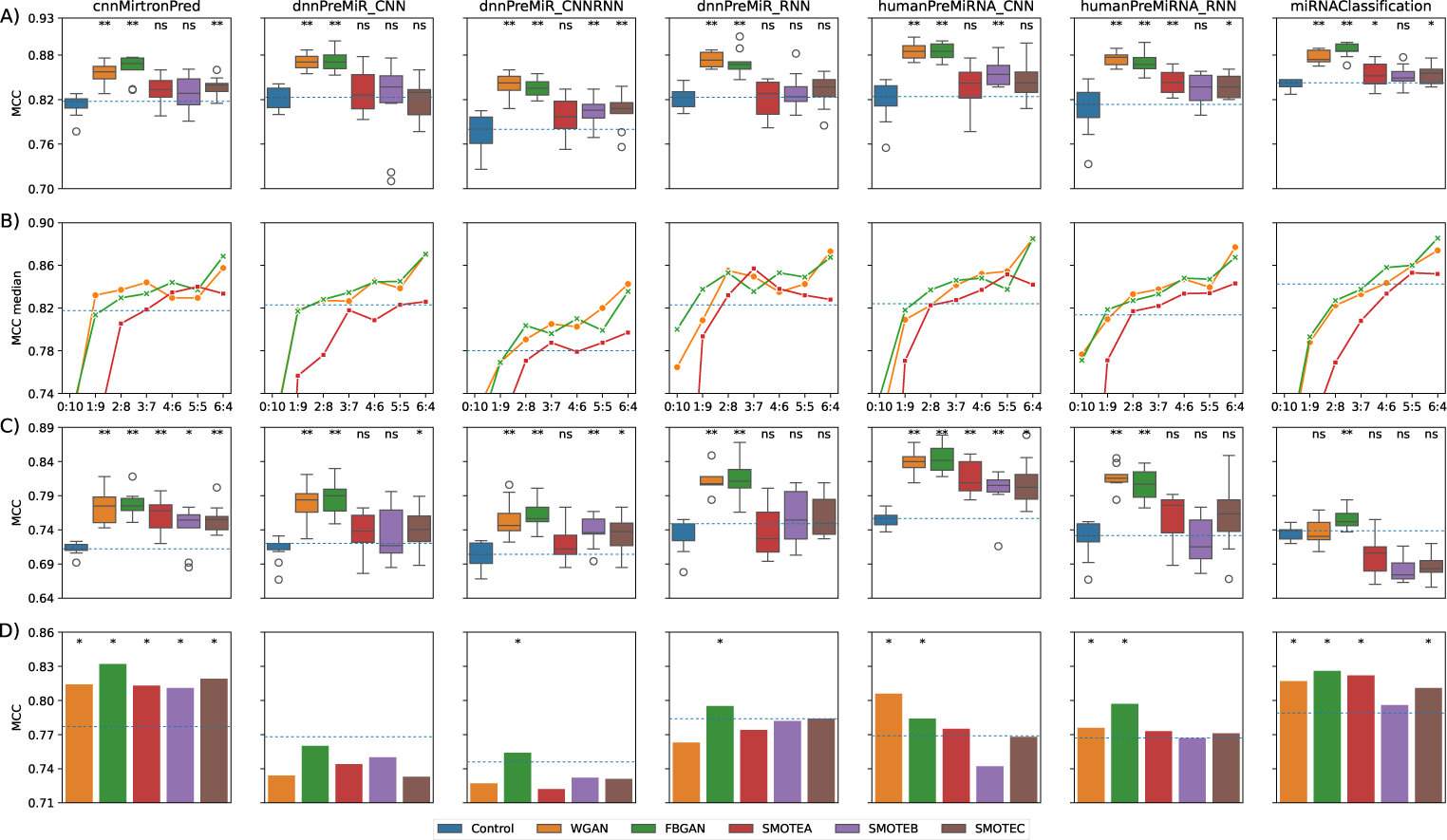
**A) Experiment I - Boxplot for the MCC comparison between control group and experimental groups for the classification tools.** Each boxplot comprises 10 data points (which is the number of training/test of the classification tool for each group). The asterisks (* and **) represent significance at *p <* 0.05 and *p <* 0.01, respectively, and ns = not significant; **B) Experiment II - MCC median for different ratios between real samples and synthetic samples.** Each point in the chart represents a median of 10 Matthews Correlation Coefficient (MCC). The 6:4 ratio represents the obtained median in the Experiment I, and the blue horizontal line shows the median of the control group; **C) Experiment III - Boxplot for the MCC comparison between control group and experimental groups for the classification tools**. Each boxplot comprises 10 data points (which is the number of training/test of the classification tool for each group). The asterisks (* and **) represent significance at *p <* 0.05 and *p <* 0.01, respectively, and ns = not significant; **D) Experiment IV using Dataset IV (M. musculus).** Each subplot corresponds to a different classification tool, and the coloured bars represent the MCC of the classification by the corresponding model of the experimental groups. The asterisk (*) denotes an improvement ≥ 1% compared to the control experiment.

The addition of GAN-based synthetic data improved performance across all tools compared to the control group, with a statistically significant difference (*p <* 0.01). The average difference between Matthews correlation coefficient (MCC) of the control group and the experimental groups WGAN-GP and FBGAN was 0.377 and 0.384, respectively. This demonstrates that both WGAN and FBGAN effectively extracted and learned the relevant features for classification between canonical miRNAs and mirtrons. Consequently, they generated mirtron sequences with the essential features to improve the classification tools.

Regarding the SMOTE experimental groups, the addition of SMOTE-based synthetic data improves the performance compared to the control group. However, this improvement was statistically significant in only 8/21 of the results (*p <* 0.05). The average difference between the control group and the experimental SMOTE groups ranged from 0.119 to 0.133. This suggests that while balancing with synthetic data using the SMOTE method can enhance classification, its effectiveness may vary depending on the context and classification tool used.

The exploratory analysis of the training sets shows a clear separation between the two classes of real samples (canonical miRNAs and mirtrons). The synthetic mirtron samples generated by FBGAN exhibit significant overlap with the real mirtron samples (Fig. 2A). In contrast, the synthetic samples produced by SMOTEA do not show the same level of overlap, although they are clustered near the real mirtron samples (Fig. 2B).

**Fig. 2.**
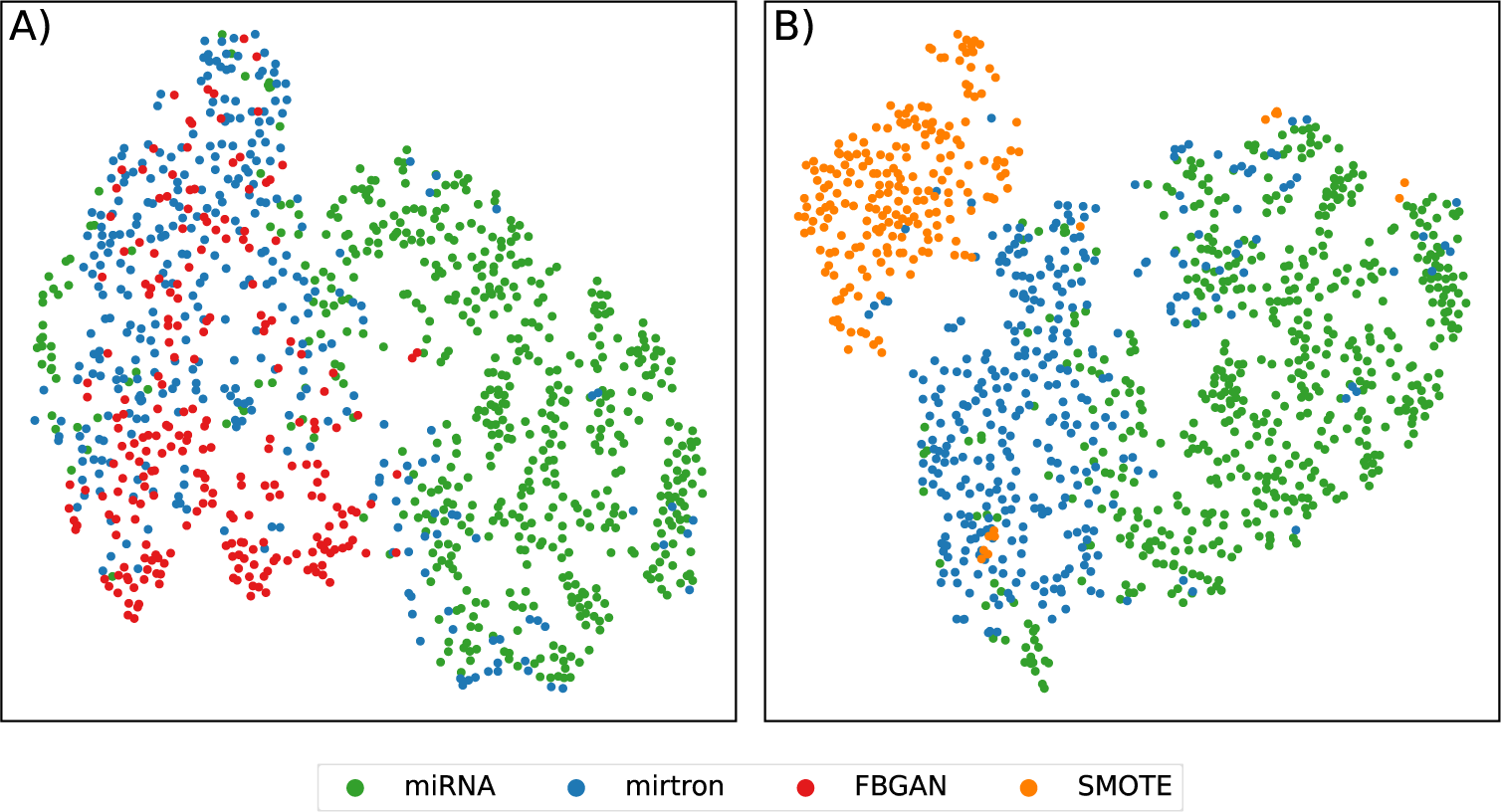
t-SNE analysis of training sets for FBGAN and SMOTEA experimental groups. The **A** plot shows the distribution of the FBGAN experimental group, while **B** plot shows the SMOTEA group. The miRNA and mirtron categories are represented by green and blue points, respectively, and the synthetic data points are represented in red (**A**) and orange (**B**).

The exploratory analysis of the test sets revealed that misclassified samples are predominantly located in regions with intersection between the two classes of samples (Fig. 3). However, when compared to the experimental FBGAN group, it is evident that there is a significant reduction in the number of incorrectly classified samples (from 56 large points in control group to 26 in FBGAN experimental group), especially in the intersection region of the data. Of the 9 mirtron samples misclassified in the experimental FBGAN group, 4 samples (hsa-mir-5004, hsa-mir-6891, hsa-mir-6740, hsa-mir-6839) are located in a region with a higher density of miRNAs, and 1 sample (hsa-mir-6853) is located in the intersection region. Of the 17 incorrectly classified miRNA samples, 4 samples (hsa-mir-7159, hsa-mir-6770-1, hsa-mir-4638, hsa-mir-6859-3) are located in a region with a higher density of mirtrons, and 5 samples (hsa-mir-4695, hsa-let-7e, hsa-mir-4800, hsa-mir-4700, hsa-mir-7107) are located in the intersection region. These misclassifications may be due to the inherent overlap in the feature space between different classes, making it difficult for the model to distinguish them in such regions.

**Fig. 3.**
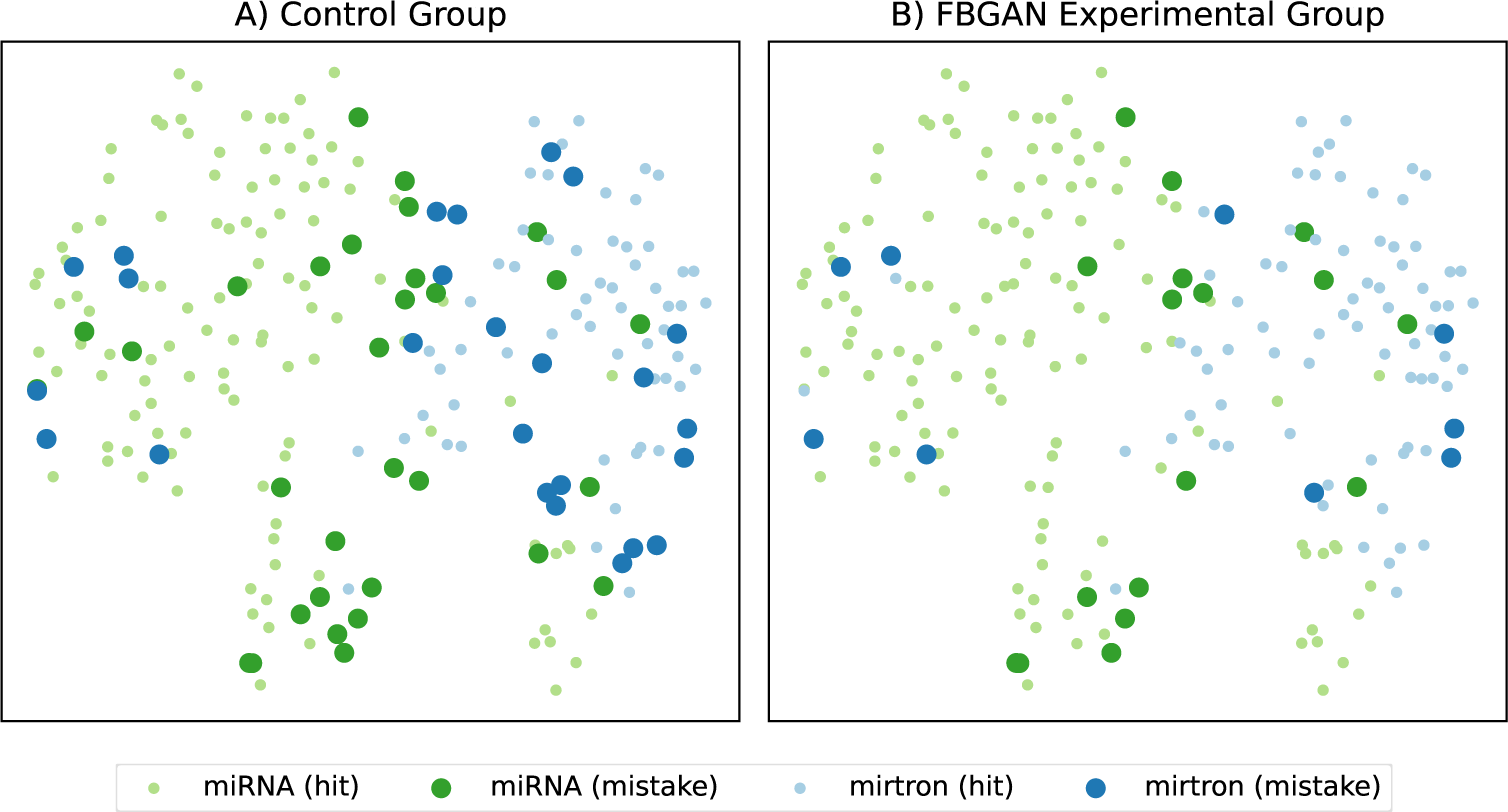
t-SNE analysis of test set and the classification results of all tools. Both the plots show the two-dimensional space of test set calculated by t-SNE. The green and blue points represent the miRNA and mirtrons data, and the big points represent the misclassified samples by the tools. The **A** plot represents the results based on control group, while the **B** plot shows the results based on FBGAN experimental group.

These analyses were conducted using t-SNE. Consistent findings were observed when repeating these analyses using Uniform Manifold Approximation and Projection (UMAP) [34] and Pairwise Controlled Manifold Approximation (PaCMAP) [35] (Supplementary Fig. S3-S6).

### 2.2 Impact of the Ratio between Real and Synthetic Data on Classification Performance

The objective of the Experiment II was to evaluate if the datasets created in the Experiment I could optimize classification tools in scenarios where the training set presented an imbalance of up to 100%. Here, six hypothetical scenarios were elaborated with distinct ratios between real and synthetic data. Fig. 1B summarizes the results of this experiment.

In general, the combination of real and synthetic data brought benefits compared to the control group. Except for the 0:10, 1:9 ratios and the results of miRNAClassification tool, all combinations with data generated by methods based on GAN showed improvements. With SMOTE, the results vary considerably among the classification tools, with the best improvements achieved for cnnMirtronPred and HumanPreMiRNA RNN.

These results corroborate the Experiment I and help answer the question: What is the minimum necessary proportion of real data to train a classification tool? The results with synthetic data generated by GAN showed that it was possible to improve classification tools using at least 20% of real data (or up to 80% of synthetic data) for the minority class. No improvement or irrelevant improvements were observed in scenarios using less than 20% real data.

### 2.3 Only half of the positive data

In the Experiment III, the number of positive samples (mirtrons) were reduced to generate new dataset (Supplementary Table S5), allowing us to evaluate the effectiveness of our generative approaches in scenarios where the number of real samples is more restricted. This reduction in real samples led to some consequences for this scenario: (i) generative approaches had less data to create new data; (ii) the training set for the control group became more imbalanced; (iii) it was necessary to use more synthetic data to balance the training set in the experimental groups.

The results showed that the addition of GAN-based synthetic data improved the performance of almost all tools compared to the control group (Fig. 1C). The only exception was the miRNAClassification tool using synthetic data from WGAN-GP, where the results showed a reduction in the median Matthews correlation coefficient (MCC).

Regarding the SMOTE experimental groups, the addition of SMOTE-based synthetic data was beneficial in 15/21 cases, of which 9 were statistically significant (*p <* 0.05). The other results, especially miRNAClassification, showed that adding SMOTE-based synthetic data did not improve the tool. Conversely, the SMOTE interpolation mechanism showed decreased performance with a smaller dataset.

These findings support the results of the Experiment I, showing that GAN approaches could extract and learn the relevant features for classification between canonical miRNAs and mirtrons, even with a smaller amount of mirtrons samples for the training. Meanwhile, the interpolation mechanism of the SMOTE has been shown to lose performance when there is a smaller amount of data.

### 2.4 Generalization Evaluation of Models Trained with Synthetic Data

In the Experiment IV, the generalization capability of the trained models in the Experiment I was investigated. In other words, these models were trained on Dataset I (*H. sapiens*) and evaluated in another species. Fig. 1D shows the classification results for Dataset IV (*M. musculus*). Additionally, Supplementary Fig. S7 and S8 show the results for Datasets II (*H. sapiens*) and III (*M. mulatta*), respectively.

The cnnMirtronPred tool achieved improvements in all analyses, followed by the miRNAClassification tool, which showed improvements in four analyses. The other five tools showed improvements in 6 out of 15 analyses, with improvements observed only when trained with WGAN-GP and FBGAN data.

Considering the results in the three datasets, the cnnMirtronPred tool’s models were the only ones that showed improvements in generalization ability in all analyses. The dnnPreMiR RNN tool showed an increase in generalization in 11 analyses, and the HumanPreMiRNA RNN and miRNAClassification tools showed improvements in 10 analyses each. However, when summarizing the gains and losses, the cnnMirtronPred and miRNAClassification tools showed the highest overall increases in MCC, with 1.543 and 0.678, respectively. In contrast, only the dnnPreMiR CNN and dnnPreMiR CNN RNN tools showed a reduction in generalization ability (MCC decrease of 0.257 and 0.086, respectively).

Interestingly, cnnMirtronPred and miRNAClassification tools are specifically designed for the classification of mirtrons and canonical miRNAs, which could explain their higher increases in MCC when classifying data from other species (and from different human origins). The other five tools were developed for miRNA identification, not specifically for distinguishing canonical miRNAs from mirtrons.

Regarding the generative approaches, FBGAN was the most effective, improving classification in 14 out of 21 analyses, followed by WGAN-GP and SMOTEC with 13 and 12 improvements, respectively. The SMOTEA and SMOTEB approaches also contributed, improving results in 11 out of 21 analyses.

Analyzing the classification tool architectures and the types of generative approaches, the tools based on convolutional neural network (CNN) architectures benefited the most from synthetic data created by our GANs.

In summary, cnnMirtronPred was the tool that most increased generalization ability compared to the control group, while dnnPreMiR CNN had the lowest gain. The most effective generative approach was FBGAN, demonstrating the greatest increases in generalization of the seven tools evaluated when trained with its synthetic data.

## 3 Discussion

This work proposes GENNUS applied to enhance the mirtron classification sequence. Two GAN-based methods were developed to generate sequences along with three SMOTE-based approaches for balancing nucleotide-level datasets. Our research shows that incorporating synthetic data generated by GANs can greatly improve the performance of classification tools, particularly in scenarios characterized by limited real data availability. Also, the advantage of these proposed methods is that they do not require feature engineering.

The combination of real data and the data created by our approaches can significantly improve the training of new models. This capability can be crucial in scenarios with limited real data, offering a significant advantage in the development and optimization of advanced machine learning models.

The experimental groups using GANs outperformed those using SMOTE. We hypothesize that GANs are more effective at capturing and reproducing the complexity and inherent diversity of real data. Unlike SMOTE, which generates new examples by simple linear interpolation between existing data points and often produces similar synthetic examples, GANs employ an adversarial process between a generator and discriminator. This process allows GANs to create more varied and realistic synthetic examples that more accurately reflect the true data distribution. As a result, models trained on data balanced by GANs benefit from a richer and more diverse representation of the minority class, enhancing their generalization and classification performance.

Investigation of features in a two-dimensional space allowed us to analyze the distribution and separation of data from various perspectives: by class (mirtrons and miRNAs), by classification performed by the tools (correct or incorrect), and by classification performed in the control and experimental groups (before and after combining synthetic data for training the tools). This allowed us to identify that the data synthesized by GAN approaches have a large overlap with the real data, while the synthetic data based on SMOTE has little overlap.

Experiments varying the proportions of real and synthetic data revealed valuable insights into the balance required for optimal model performance. Specifically, it was found that the use of at least 20% of real data in combination with synthetic samples provides a significant increase in the classification precision. These findings suggest that, while synthetic data can improve model performance, a minimum proportion of real data is necessary to maintain robust generalization.

To conclude, the use of synthetic data can be an effective strategy to improve the performance of neural networks, especially when there are limitations in the available real data. However, it is crucial to ensure that the synthetic data is representative and complements the real data in a beneficial way. Continuous evaluation and experimentation with different methods of synthetic data generation are fundamental to achieving proper balance and maximizing the benefit of this approach.

This study raises two critical questions that remain unanswered. First, can these results be replicated in models with different architectures, such as Transformers, which use tokenization processes for handling input data? Second, is it possible to generalize these findings to other biological problems? More research and additional studies are required to address these questions and extend the applicability of these findings.

## 4 Methods

### 4.1 Datasets

Real samples from *Homo sapiens*, *Mus musculus*, and *Macaca mulatta* were obtained from the mirtronDB database [36] and the miRBase database [37]. The mirtronDB specifically focuses on mirtrons in chordates, vertebrates, and plants, while miRbase gathers and accepts miRNA entries from researchers and publications across various organisms. Additionally, the Wen Dataset [3], previously utilized by cnnMirtronPred [9] and miRNAClassification [12], was also collected.

The collected real samples were divided into four distinct sets comprising mirtrons and canonical miRNAs (Supplementary Table S2). Dataset I consisted exclusively of samples from the Wen Dataset (*Homo sapiens*). Datasets II, III, and IV were created by merging samples collected from mirtronDB and miRBase, with these three sets comprising samples from *Homo sapiens*, *Macaca mulatta* and *Mus musculus*, respectively.

To ensure uniqueness, intersections between mirtron sequences from mirtronDB and miRBase were identified using BLAST (blastn v2.14.1 with a minimum of 90% of identical positions [38]), and overlapping samples from miRBase were removed. Furthermore, samples already present in Dataset I were removed from Dataset II.

### 4.2 miRNAs and mirtrons classification tools

From the literature, seven classification tools were selected to be used in the experiments to evaluate the data augmentation approaches (Supplementary Table S1). To exclusively investigate the effect of changes in the training datasets on the performance of classification models, all seven selected tools were re-trained using their default settings (according to their respective repositories). This procedure ensures that the analysis focuses on the impact of the different datasets used for training, avoiding the influence of hyperparameter tuning or specific optimizations for each tool.

By keeping the parameters constant, we ensure that any variation in performance is solely attributable to the characteristics of the training datasets used.

The cnnMirtronPred [9] and miRNAClassification [12] tools were developed to identify and differentiate the intrinsic differences between canonical miRNAs and mirtrons. These tools were trained and tested using the Wen Dataset [3]. On the other hand, the dnnPreMiR [10] and HumanPreMiRNA [11] tools were developed to detect precursor miRNA samples by recognizing the features of secondary hairpin structures and identifying potential miRNAs. For training, precursor miRNA samples served as the positive set, while RNA samples from coding regions were used to create the negative set.

About the architecture, cnnMirtronPred, miRNAClassification, HumanPreMiRNA CNN and dnnPreMiR CNN are based exclusively on Convolutional Neural Networks (CNN), and HumanPreMiRNA RNN and dnnPreMiR RNN tools are based on Recurrent Neural Networks (RNN). dnnPreMiR CNN RNN tool uses a hybrid architecture with CNN layers for feature extraction from input data and RNN layers for sequence prediction.

### 4.3 Data Augmentation by Generative Adversarial Networks

The classical architecture of a GAN [39] consists of two main networks: a generator *G* and a discriminator *D* (Fig. 4). The generator *G* transforms an input noise *z* into synthetic data *G*(*z*). The discriminator *D* receives the input data *x* (real and synthetic) and produces an output *D*(*x*), aiming to distinguish between real and fake data. The output *D*(*x*) is limited to [0, 1], representing the probability that the data *x* is real. The goal of the generator *G* is to generate data points so convincingly that the discriminator *D* cannot distinguish them from real data.

**Fig. 4.**
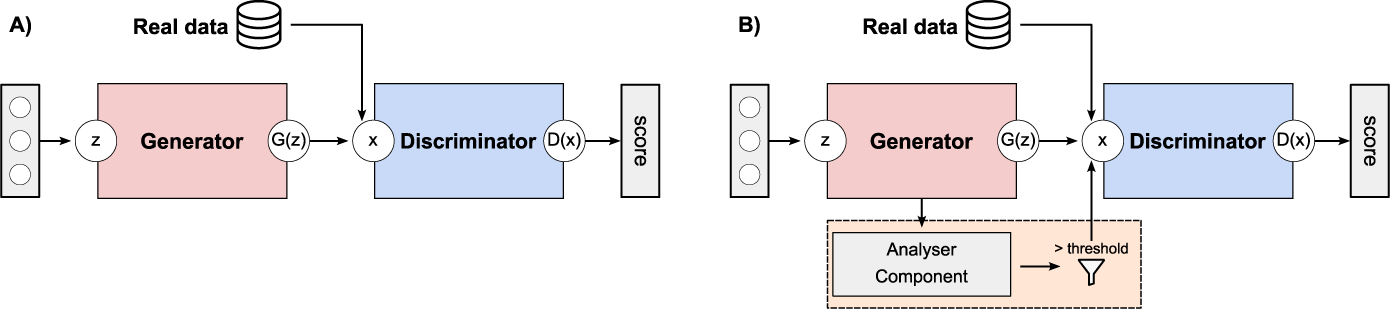
Basic schematic of a (A) WGAN-GP and a (B) FBGAN architecture. The generator *G* and discriminator *D* are both parameterized as deep neural networks. The real data is encoded using one-hot vectors, and the generator learns a continuous approximation to this encoding. In a FBGAN (B) architecture, the feedback-loop mechanism consists of two components: a WGAN-GP architecture and the Analyser Component. The Analyser Component scores each generated data, and data above of the threshold are added to the training set.

Training proceeds by alternating between training phases for the discriminator *D* and the generator *G*. The discriminator is trained using both real data (*P_real_*) and synthetic data (*P_z_*). Real data is sampled from a dataset containing real data points. In contrast, synthetic data is obtained by sampling a latent variable *z*, and passing it through the generator *G* to produce synthetic data *G*(*z*). During training, the discriminator’s objective is to correctly classify whether the input data *x* is real or synthetic without knowing its origin. This training process involves multiple iterations where the parameters of both the discriminator *D* and the generator *G* are adjusted to improve the quality of the generated data, making it more closely approximate the distribution of real data.

In this study, the Wasserstein GAN with Gradient Penalty (WGAN-GP) and the Feedback GAN (FBGAN) were specifically used. These variations introduce different mechanisms and objectives to enhance training stability and improve the quality of generated samples.

#### 4.3.1 Wasserstein GANs with Gradient Penalty

The Wasserstein GANs with Gradient Penalty (WGAN-GP) [40, 41] was proposed to address some issues of traditional GANs [39], such as mode collapse and training instability. The central concept of a WGAN-GP is to use the Wasserstein distance to measure the difference between the generated data distribution *P_z_* and the real data distribution *P_real_*. In a WGAN-GP, the discriminator’s output *D*(*x*) is no longer constrained to [0, 1]. Instead of predicting the probability of *x* being real or fake, the discriminator provides scores related to the realism of *x* [40].

The components of WGAN-GP of this work were determined empirically (Supplementary Fig. S9 and Supplementary Tables S7-S10). Both the discriminator *D* and the generator *G* employ five chained residual blocks (ResBlock - Supplementary Fig. S9C). Each ResBlock consists of two Rectified Linear Unit (ReLU) layers followed by a Conv1D layer, and the output of last layer is multiplied by *r* (*r ≤* 1) and added to the ResBlock input. The Conv1D layers have a padding size equal to 2.

The generator *G* (Supplementary Fig. S9A) is composed of five sequential layers: a Linear layer, a Reshape layer, five chained ResBlock, a Conv1D layer, and a Softmax layer. The input *z* is a numeric vector of length 100, and the output *G*(*z*) is a matrix of dimensions 121 × 5. An *argmax* function is applied to *G*(*z*) to generate a single nucleotide at each position.

The discriminator *D* (Supplementary Fig. S9B) consists of four sequential layers: a Conv1D layer, five chained ResBlock, a Reshape layer, and a Linear layer. The input *x* is a numeric vector of dimensions 121 × 5, and the output *D*(*x*) is a score indicating the realism of *x*.

Mirtrons sequences from Dataset I were used as input to train the WGAN-GP. In a pre-processing step, the lengths of all RNA sequences were standardized by padding each with a varying number of ‘P’ nucleotides at the end, extending them to a maximum length of 121 nucleotides (this length represents ≥ 95% of our mirtron samples). Following this, one-hot encoding was employed to represent each base in the samples (*P* = [1, 0, 0, 0, 0], *A* = [0, 1, 0, 0, 0], *T/U* = [0, 0, 1, 0, 0], *G* = [0, 0, 0, 1, 0], and *C* = [0, 0, 0, 0, 1]). Consequently, each base was transformed into a five-dimensional vector, resulting in the vectorization of each mirtron sequence into a (121,5)-dimensional vector.

The WGAN-GP was trained for 100 epochs, and the batch size was set to 32. The ratio 5 : 1 between the discriminator and generator training steps was selected. To optimize the network, the Adam algorithm optimizer [42] with *β*_1_ set to 0.5 and *β*_2_ set to 0.9 was employed. The learning rates for the generator and discriminator were set to *γ* = 10^−4^.

The discriminator was trained using both real and synthetic data. The real data was taken from one-hot encoded mirtrons dataset. The synthetic data was obtained by sampling the latent variable *z* with a normal distribution and asking the generator *G* to transform the random noise *z* into synthetic data *G*(*z*). For the generator training, fake data samples were generated and passed to the discriminator to be scored.

#### 4.3.2 Feedback GAN

The Feedback GAN (FBGAN - Fig. 4B) [27] was proposed as a novel feedback-loop architecture to optimize the synthetic gene sequences for desired properties. The feedback-loop mechanism consists of two components: a GAN, which follows the WGAN-GP architecture, and the Analyser. The Analyser takes a sample and assigns a ‘favourability’ score to the sequence. The GAN and Analyser are linked by the feedback mechanism. At each epoch, several samples are obtained from the generator and input into the Analyser. The Analyser assigns a score to each sample, and all samples with a score above the threshold are input back into the discriminator’s dataset (here with *threshold >* 0.95). Only the last *nlast* generated samples are kept from the discriminator’s training dataset, and the oldest generated are removed. The GAN is then trained as usual for each epoch. As the feedback process continues, all the generated samples of the discriminator training set are replaced repeatedly by high-scoring generated sequences.

In this work, a FBGAN based on WGAN-GP was implemented, leveraging the generator *G* and discriminator *D* components. The Analyzer component was designed following the neural network architecture of the WGAN-GP discriminator (Supplementary Fig. S9B). A Sigmoid layer was added to the output layer, allowing the neural network to produce values between 0 and 1, representing the ‘favourability’ of the sample. The training dataset (Section 4.5.1 - Preparing the data (B)) was used as input to train the Analyzer Component for 100 epochs, and the batch size was set to 32. To optimize the network, the Adam algorithm optimizer with *β*_1_ set to 0.9, *β*_2_ set to 0.999, and learning rate set to *γ* = 10^−4^ was employed. Based on the test dataset, the trained Analyzer Component’s accuracy is 0.938. After that, the training process for the FBGAN followed the same procedure as described for the WGAN-GP.

### 4.4 Data Augmentation by SMOTE

Synthetic Minority Over-sampling TEchnique (SMOTE) [43] is a pre-processing technique used to address an imbalanced dataset. To the best of our knowledge, there is no variation of SMOTE for primary sequences of nucleotides, so three SMOTE-based steps were followed to generate nucleotide sequences.

Firstly, the length of the training set was normalized to 121 nucleotides (this length represents ≥ 95% of our mirtron samples). Sequences longer than this length were truncated, and shorter sequences were padded with the character ‘P’, both at 3’ end (right side). Next, the nucleotides were converted to their corresponding numeric values (*P* = 0, *A* = 1, *C* = 2, *G* = 3, *T* = 4). These transformations are prerequisites for applying SMOTE, as its input must be a two-dimensional numeric matrix. This process resulted in a numeric matrix of dimensions *L ×* 121, where *L* is the size of the dataset.

After this, an iterative process was applied to generate three synthetic datasets of nucleotide sequences (SMOTEA, SMOTEB, and SMOTEC). First, SMOTE was applied to the numeric matrix to create synthetic samples. For each generated sample, any padding nucleotides ‘P’ were removed from the end of the sequence. Next, in the SMOTEA set, the samples that still contained ‘P’ nucleotides were removed. In the SMOTEB set, ‘P’ nucleotides were replaced with another randomly chosen nucleotide. In the SMOTEC set, the ‘P’ nucleotides were removed. This process was repeated iteratively until each dataset reached 3,000 samples.

### 4.5 Experiment designs

In the experimental design section, four key experiments were conducted to evaluate the impact of synthetic data on ncRNA classification. First, the balance between real and synthetic data was assessed (Experiment I). Next, the effect of varying real:synthetic data ratios on model performance was analyzed (Experiment II). In Experiment III, the effectiveness of generative approaches was tested when real data was reduced by half, increasing class imbalance. Finally, the generalization of models trained on human data to other species, including mice and *Macaca mulatta*, was evaluated (Experiment IV).

#### 4.5.1 Experiment Design I: Evaluating Balance with Synthetic Data

The objective of this experiment was to evaluate whether synthetic data generated by our proposed approaches can enhance the performance of classification tools. The experimental design is illustrated in Fig. 5, with the tasks described as follows:

**Fig. 5.**
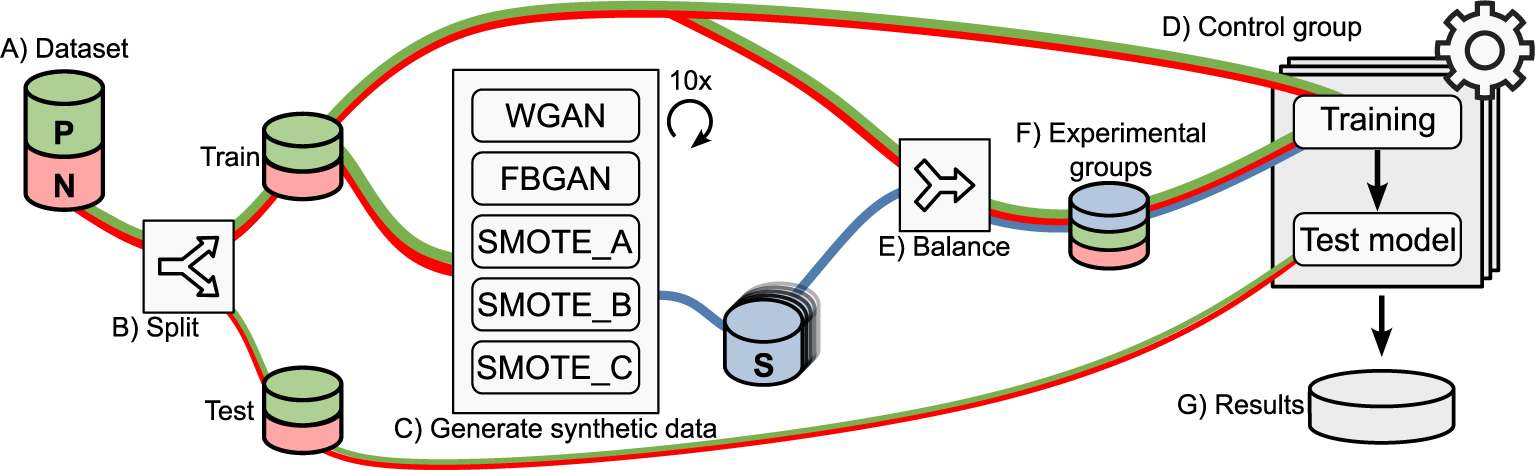
The Process of Generating and Evaluating Synthetic Data. (A) Real-world data is collected and (B) split into training and test sets. (C) GenAI approaches generate synthetic data, which (E) is used to balance the training set. The classification tools were trained using (D) real data and (F) balanced datasets. Finally, (G) the models are tested using the test set.

**Dataset (Task A - Fig. 5A):** The Dataset I was used for this experiment, which consists of 417 mirtrons as positive samples and 707 canonical miRNAs as negative samples.

**Preparing the data (Task B - Fig. 5B):** The Dataset I was split using a stratified approach, with 80% of the data randomly selected for training, while the remaining 20% was set aside for testing. The distribution of training and test sets is shown in Fig. 5B and Supplementary Table S3.

**Generating synthetic data (Task C - Fig. 5C):** WGAN-GP and FBGAN were trained ten times, generating ten different versions of the Generator (as described in Section 4.3). For each Generator, 3,000 synthetic samples were produced. Furthermore, ten synthetic datasets were created for each of the three SMOTE approaches, each containing 3,000 samples (as detailed in Section 4.4). In total, 50 synthetic datasets were generated (10 × 5 approaches, summarized in Supplementary Table S6).

**Control group (Task D - Fig. 5D):** The control group results were used as a baseline for comparison with the experimental groups using synthetic data. Each of the seven classification tools (listed in Supplementary Table S1) was trained 10 times using only real data, and all models were evaluated on the same test set.

**Balancing the training dataset (Task E - Fig. 5E):** The balancing process uses synthetic data to balance the number of samples in a two-class set. It takes an unbalanced training set and a synthetic dataset as input, calculates the class imbalance, and adds synthetic samples to the minority class. The result is a new balanced set with synthetic data combined into the minority class - the samples of mirtrons. The original input sets remain unchanged.

This process was applied to create a new balanced dataset for each of the 50 synthetic datasets created in Task C (Fig. 5C). As a result, 50 balanced training sets were obtained consisting of 565 mirtrons (334 real + 231 synthetic, or ≈ 6 : 4 ratio) and 565 canonical pre-miRNAs (Supplementary Table S3). The testing set remained unchanged. In practice, each synthetic set was used only once to balance a copy of the training set (Supplementary Table S6).

**Experimental groups (Task F - Fig. 5F):** The 50 balanced training sets - created in Task E - were categorized by generative approach in five experimental groups (WGAN, FBGAN, SMOTEA, SMOTEB and SMOTEC). Each group contain 10 datasets balanced by the correspondent generative approach.

Subsequently, each of the seven classification tools (Supplementary Table S1) were trained once for each of the 50 training sets, resulting in 50 models for each tool. Finally, all trained models were tested with the same test set.

**Analyzing results (Task G - Fig. 5G)**: A total of 420 results were obtained using the same test set. These results were categorized by classification tool and group (the control or one of the five experimental groups). In other words, were obtained 70 results for the control group (Task D - Fig. 5D) and 70 results for each experimental group (10 balanced datasets x 7 tools).

The Paired Wilcoxon-Signed Rank Test was used to compare statistically the MCC values between the control and experimental groups, and the significance levels were considered as *p ≤* 0.05 and *p ≤* 0.01 for all statistical analysis. The wilcoxon function of scipy.stats package (version 1.11.4) was used to compute all tests [44].

Additionally, for this experiment, two exploratory analyses were performed. Firstly, a feature extraction step was employed to extract feature vectors based on Minimum Free Energy (MFE), sample length, GC content, and the count of nucleotides A, C, G, and T for each sample of the training and test sets.

In the first analysis, the training datasets of experimental groups were examined. T-distributed Stochastic Neighbor Embedding (t-SNE) [45] was used to reduce the feature vectors to 2D vectors and visualize the data separation in balanced training sets. These features were passed as input into a t-SNE reducer object (using default parameters, *n iter* = 2500 and *random state* = 42) from the sklearn python package.

In the second analysis, the test results were evaluated by comparing the control group with the experimental group that showed the best performance (FBGAN). Two groups were defined: Hits and Mistakes. The Hits group consists of samples correctly classified by all seven tools, while the Mistakes group includes sequences that were incorrectly classified by at least one of the tools. T-distributed Stochastic Neighbor Embedding (t-SNE) [45] was used to reduce the feature vectors.

#### 4.5.2 Experiment Design II: Evaluating the Effect of Real vs. Synthetic Data Ratios

The goal of the Experiment II was to build on the findings from Experiment I by varying the proportion of synthetic data in the training set, ranging from 40% (6:4 ratio of real:synthetic) to 100% synthetic data. This allowed to assess the minimum real data required to maintain strong classification performance across the tools. To achieve this, the experimental setup from Experiment I was replicated, with modifications to the training set balancing process (Task E, detailed in Section 4.5). In Experiment I, the positive class (mirtrons) consisted of 565 samples, with a ratio of approximately 6:4, including 334 real and 231 synthetic samples. The negative class (canonical miRNAs) also had 565 real samples, resulting in a balanced dataset with equal representation of both classes. In Experiment II, an additional parameter, the real:synthetic ratio, was introduced to control the proportion of real and synthetic samples in the minority class. This parameter adjusted the number of real and synthetic samples used to compose the mirtron class, resulting in new balanced datasets based on the specified ratio.

For this Experiment, the focus was exclusively on the synthetic data created by the WGAN-GP and FBGAN approaches, as they achieved the best results in the Experiment I. Additionally, SMOTEA was used to represent SMOTE-based approaches. The following real:synthetic ratios were tested for the mirtron class: 5:5 (50% real, 50% synthetic), 4:6, 3:7, 2:8, 1:9, and 0:10 (100% synthetic). The balancing process was applied for each generative approach and ratio, resulting in 180 new datasets, categorized into 18 experimental groups based on the generative approach and data ratio (3 approaches × 6 ratios = 18 groups). The sample sizes for each set is detailed in Supplementary Table S4. Afterwards, these newly balanced training sets were used to train the seven classification tools (Supplementary Table S1), and the same testing set(from Experiment I) was used to evaluate the created models. This produced 1,260 results (180 for each tool), classified into 126 groups (each representing a specific classification tool, generative approach, and ratio). Finally, the median performance for each group was calculated, and the results were plotted for further analysis.

#### 4.5.3 Experiment Design III: Impact of Reducing Real Data with Synthetic Balancing

This experiment introduced a more challenging scenario by reducing the number of real mirtrons in the positive dataset by approximately ≈ 50%. After subsampling Dataset I, the new dataset comprised 216 stringent mirtrons as the positive class while retaining the original 707 canonical miRNAs as the negative class. This adjustment increased the imbalance between positive and negative samples to a ratio of approximately ≈ 1:3. This setup allowed for evaluating the effectiveness of the generative approaches in scenarios where real data is minimal, a common challenge in biological datasets. The design of Experiment I was replicated, using the subsampled Dataset I as input for Task B (Fig. 5B). All subsequent processes followed the same methodology as in the Experiment I. For comparison, the original sample sizes in Dataset I are listed in Supplementary Table S3, while the sample sizes for this experiment are provided in Supplementary Table S5. Finally, boxplots were calculated, and statistical tests were conducted to compare the control and experimental groups.

#### 4.5.4 Experiment Design IV: Cross-Species Evaluation of Human-Trained Models

This experiment assessed how well classification tools trained on synthetic data generalize across various species and data sources. The best-performing models from each tool and group generated in the Experiment I (Section 4.5.1) were evaluated using the Matthews Correlation Coefficient (MCC). A total of 42 models were selected (seven tools across one control group and five experimental groups). These models were then tested on Datasets II, III, and IV, which included data from *Homo sapiens*, *Macaca mulatta* and *Mus musculus*, collected from mirtronDB and miRBase (Supplementary Table S2).

## 5 Data and code availability

All data used for this research are publicly available at https://doi.org/10.5281/zenodo.13235408.

An open-source version of code to fully reproduce results has been released at https://github.com/chiquitto/GENNUS which includes the WGAN-GP, FBGAN and SMOTEs generative approaches.

## 6 Acknowledgements

The authors would like to acknowledge the support of Programa de Afastamento para Capacitação em Curso de Pés-Graduação de Docentes (Edital 076/2022) of Federal Institute of Education Science and Technology of Mato Grosso do Sul/Brasil (IFMS); CNPq (#440412/2022-6); Fundação Araućaria (Project: NAPI Bioinformatica) via grant 66.2021; and Advanced Research Computing (ARC) Team at The Rosalind Franklin Institute (UK). This study was financed in part by the Coordenação de Aperfeiçoamento de Pessoal de Ńıvel Superior - Brasil (CAPES) - Finance Code 001 [grant number PDSE-88881.933875/2024-01] in collaboration with The Rosalind Franklin Institute (UK). We thank Fabiana Rodrigues de Goes (RFI), Andre Yoshiaki Kashiwabara (UTFPR) and Douglas Silva Domingues (University of São Paulo, ESALQ/USP) for valuable comments and feedback.

## 7 Author information

Conceptualization: A.G.C and A.R.P. Experimental methodology: A.G.C, L.S.O., P.H.B. and P.T.M.S. Computacional methodology: A.G.C and M.B. Supervision: A.R.P. and R.T.R. Writing - original draft: A.G.C Final review & editing: all authors.

## 8 Ethics declarations

The authors declare no competing interests.

## 10 Supplementary information

### 10.1 Selected classification tools

**Table S1.** Tools used in evaluating DA approaches.

| Tool | Based on | Params † | Reference |
| --- | --- | --- | --- |
| cnnMirtronPred | CNN | 69,122 | [9] |
| dnnPreMiR_CNN | CNN | 60,882 | [10] |
| dnnPreMiR_RNN | RNN | 122,136 | [10] |
| dnnPreMiR_CNN_RNN | CNN+RNN | 164,488 | [10] |
| HumanPreMiRNA_CNN | CNN | 183,874 | [11] |
| HumanPreMiRNA_RNN | RNN | 253,726 | [11] |
| miRNAClassification | CNN | 170,274 | [12] |

† Trainable parameters

### 10.2 Datasets used as input

**Table S2.**
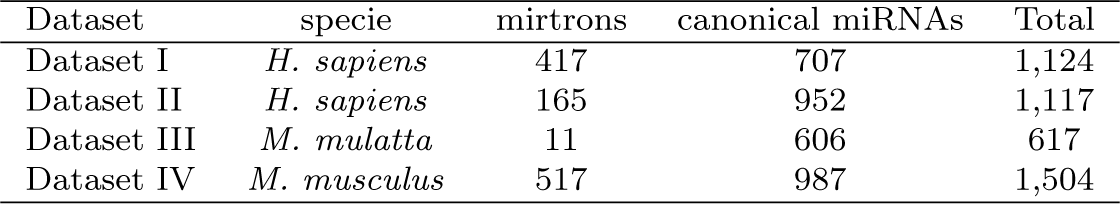
Datasets collected from mirtronDB and miRBase.

**Table S3.** Datasets used as input in Experiment I.

| Dataset | Training | Test | Total |
| --- | --- | --- | --- |
| Positive (real) | 334 | 83 | 417 |
| Positive (synthetic) <sup>1</sup> | 231 | 0 | 231 |
| Negative (real) | 565 | 142 | 707 |
| Total (real) <sup>2</sup> | 899 | 225 | 1124 |
| Total (real + synthetic) <sup>3</sup> | 1130 | 225 | 1355 |

<sup>1</sup> The synthetic samples were combined just in the training set.
<sup>2</sup> Dataset used as input in control groups
<sup>3</sup> Dataset used as input in experimental groups

**Table S4.** Ratio real:synthetic for positive samples in Experiment II.

| Dataset | real:synthetic ratio |  |  |  |  |  |
| --- | --- | --- | --- | --- | --- | --- |
|  | 0:10 | 1:9 | 2:8 | 3:7 | 4:6 | 5:5 |
| Positive (real) | 0 | 57 | 113 | 170 | 226 | 283 |
| Positive (synthetic) | 565 | 508 | 452 | 395 | 339 | 282 |
| Negative (real) | 565 | 565 | 565 | 565 | 565 | 565 |
| Total | 1,130 | 1,130 | 1,130 | 1,130 | 1,130 | 1,130 |

**Table S5.** Size of the datasets used as input in Experiment III.

| Dataset | Training | Test | Total |
| --- | --- | --- | --- |
| Positive (real) | 173 | 43 | 216 |
| Positive (synthetic) <sup>1</sup> | 392 | 0 | 392 |
| Negative (real) | 565 | 142 | 707 |
| Total (real) <sup>2</sup> | 738 | 185 | 923 |
| Total <sup>3</sup> | 1,130 | 185 | 1,315 |

<sup>1</sup> The real and synthetic samples were combined just in training set.
<sup>2</sup> Dataset used as input in control groups
<sup>3</sup> Dataset used as input in experimental groups

### 10.3 Number of datasets in the experiments

**Table S6.**
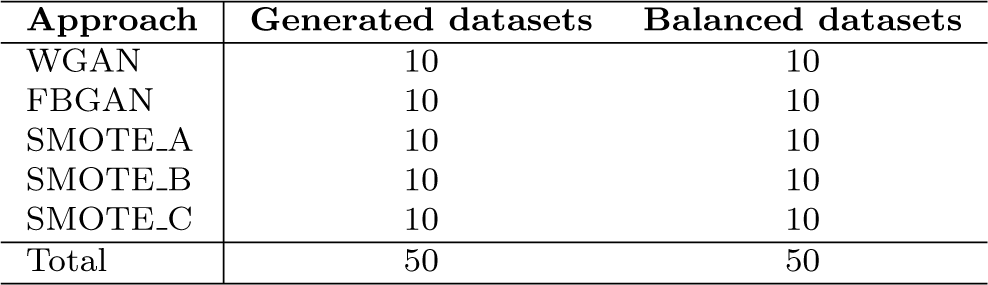
Number of datasets created by approach in the Experiment I.

| Approach | Generated datasets | Balanced datasets |
| --- | --- | --- |
| WGAN | 10 | 10 |
| FBGAN | 10 | 10 |
| SMOTE_A | 10 | 10 |
| SMOTE_B | 10 | 10 |
| SMOTE_C | 10 | 10 |
| Total | 50 | 50 |

Table S6 shows the number of datasets created by each generative approach in the Experiment I. Next, each synthetic dataset was used once to balance a copy of the training set.

### 10.4 Number of misclassified samples

Figures S1 and S2 show the upset plots for misclassified samples by the seven tools in the Experiment I - FBGAN experimental group. Intersections are displayed as a matrix. Each row corresponds to one classification tool, and bar charts on the left show the number of misclassified by the tool. Each column corresponds to a possible intersection: the filled-in circles show which set is part of an intersection, and vertical bar charts show the size of the intersection.

Just one canonical miRNA was misclassified by the seven tools (Figure S1). miRNAClassification was the best tool (four misclassified sequences), while dnnPreMiR CNN RNN was the one that presented the most errors (11 misclassified sequences).

Two mirtrons were misclassified by the seven tools (Figure S2). HumanPreMiRNA CNN was the best tool (two misclassified sequences), while dnnPreMiR RNN was the one that presented the most errors (six misclassified sequences).

**Fig. S1.**
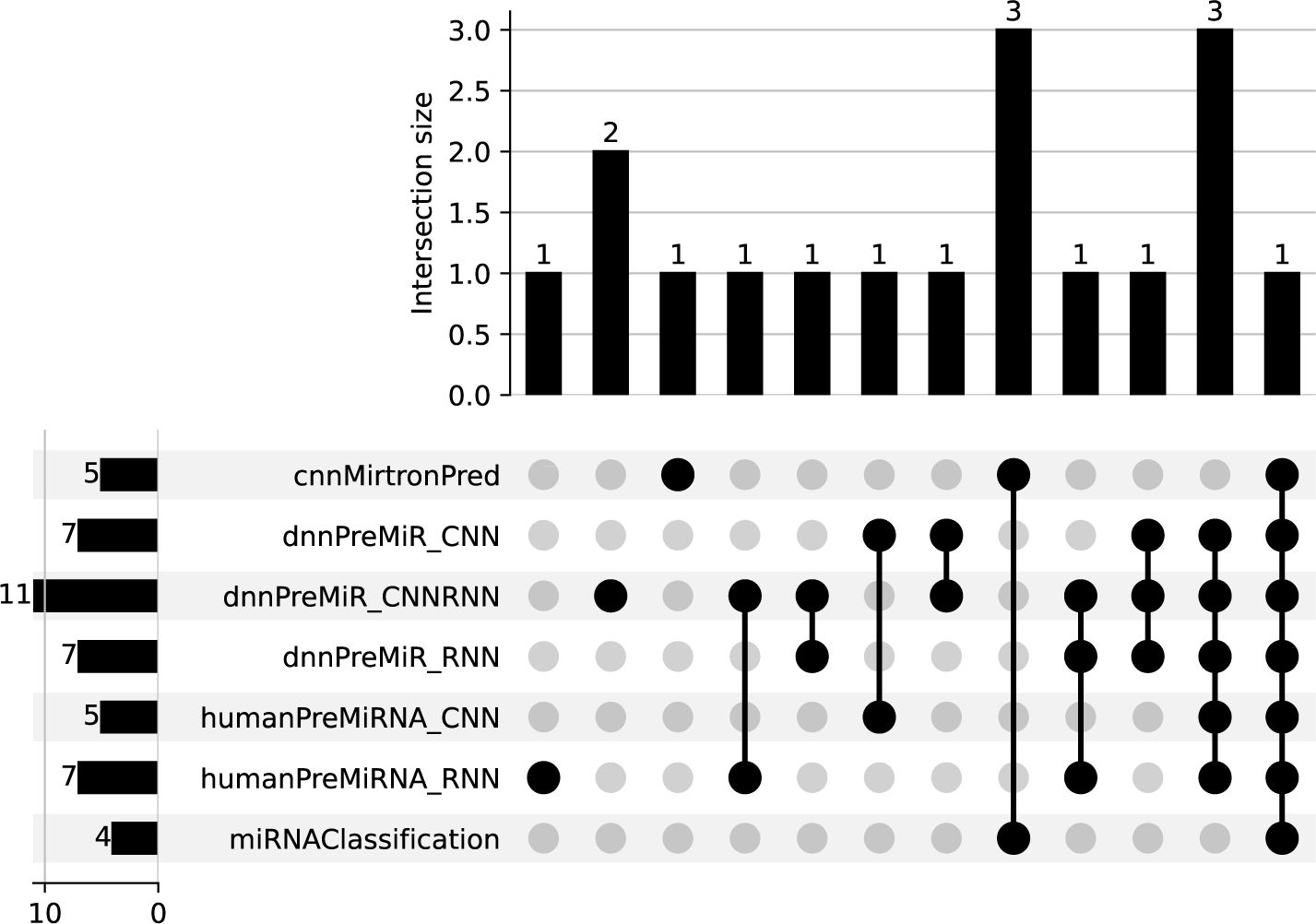
Upset plot misclassified canonical miRNAs.

**Fig. S2.**
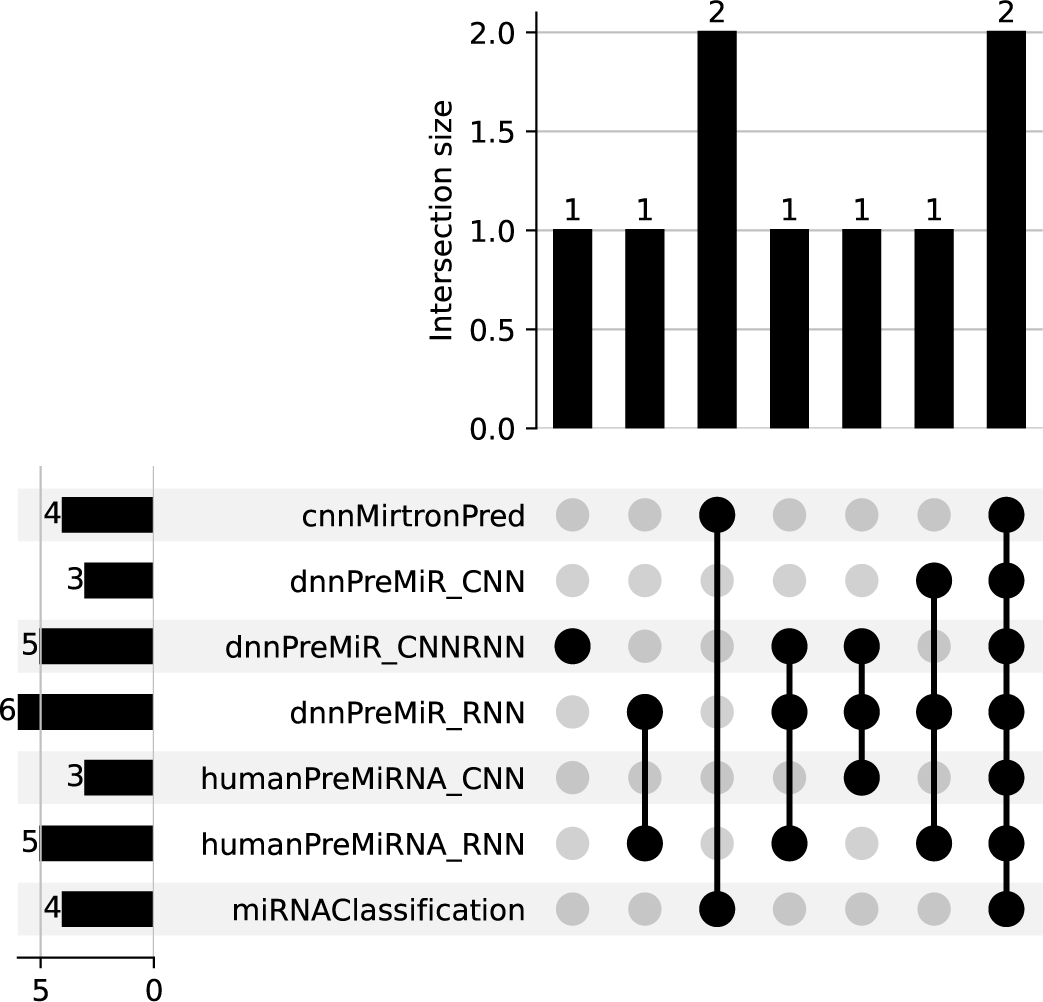
Upset plot misclassified mirtrons.

#### 10.4.1 UMAP and PaCMAP analysis of training and test sets

Uniform Manifold Approximation and Projection (UMAP) [34] and Pairwise Controlled Manifold Approximation (PaCMAP) [35] were used to reduce the features of the samples to 2D vectors and visualize the data separation. The use of UMAP and PaCMAP complements the analysis perfomed with T-distributed Stochastic Neighbor Embedding (t-SNE).

A feature extraction step was employed to extract feature vectors based on Minimum Free Energy (MFE), sample length, GC content, and the count of nucleotides A, C, G, and T for each sample of the training and test sets. These features were passed as input into a UMAP (with default parameters) from the umap-learn python package [34] and PaCMAP (with default parameters) from the pacmap python package [35]. The results are presented in Figures S3, S5 and S6.

Figure S3 shows the UMAP projections of the training set of Control Group and FBGAN experimental groups. The synthetic mirtron samples generated by FBGAN significantly overlap with the real mirtron samples (Fig. S3A). On the other hand, the synthetic samples generated by SMOTEA do not exhibit this same overlap, despite being concentrated at the real mirtron samples (Fig. S3B).

Figure S4 shows the UMAP projections of the test set. Larger points represent the 56 incorrectly classified samples in the control group (Fig. S4A) and the 26 incorrectly classified samples in the experimental FBGAN group (Fig. S4B). It can be observed that misclassified samples are predominantly located in regions with intersection between the two classes of samples. However, when compared to the experimental FBGAN group, it is evident that there is a significant reduction in the number of incorrectly classified samples, especially in the intersection region of the data. These misclassifications may be due to the inherent overlap in the feature space between different classes, making it difficult for the model to distinguish them in such regions. The same observation can be made when using PaCMAP (Figures S5 and S6).

**Fig. S3.**
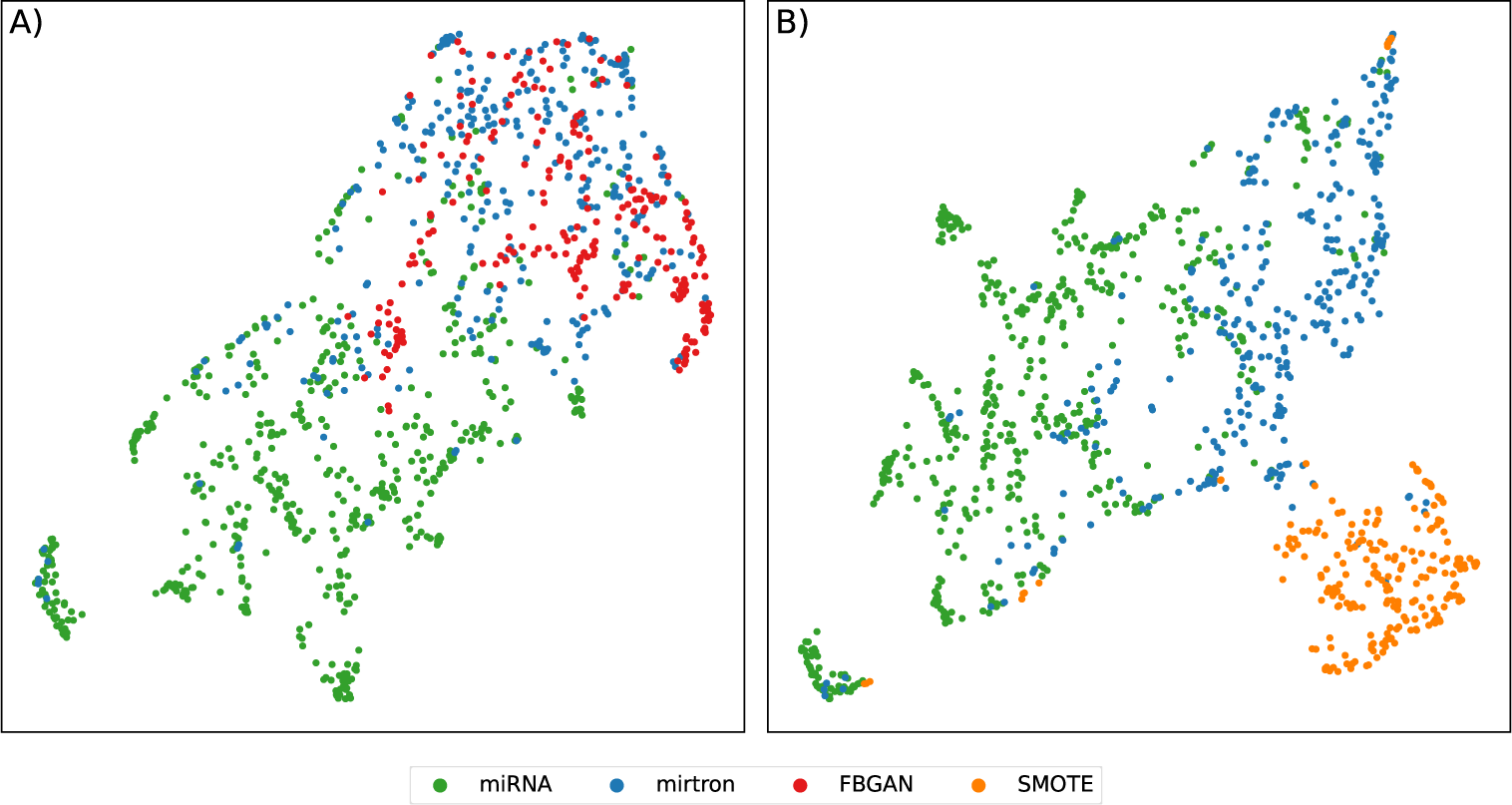
UMAP analysis of training sets for FBGAN and SMOTEA experimental groups. The **A** plot shows the distribution of the FBGAN experimental group, while the **B** plot shows the SMOTEA group. The miRNA and mirtron categories are represented by green and blue points, respectively, and the synthetic data points are represented in red (**A**) and orange (**B**).

**Fig. S4.**
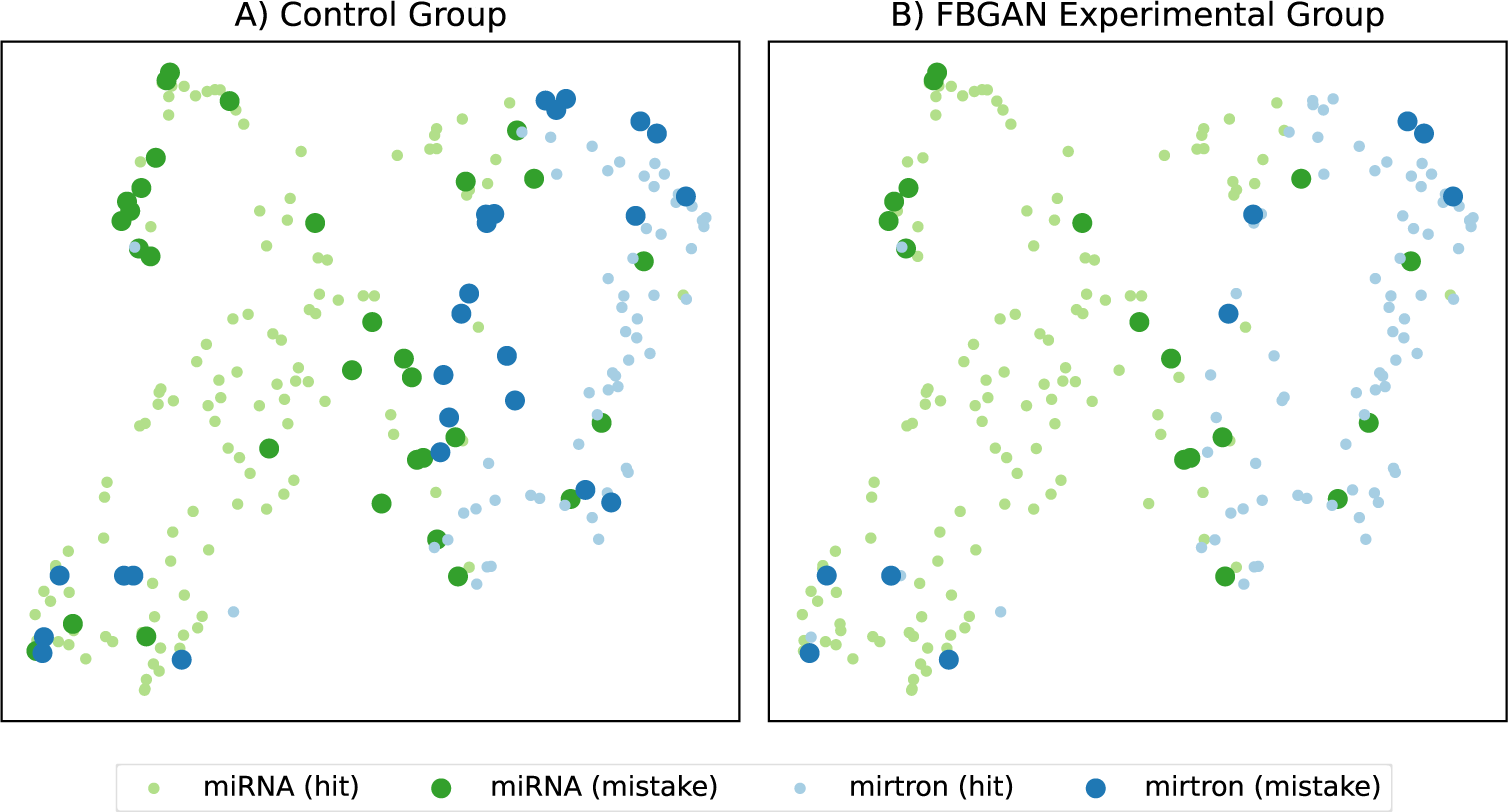
UMAP analysis of test set and the classification results of all tools. Both the plots show the two-dimensional space of test set calculated by UMAP. The green and blue points represent the miRNA and mirtrons data, and the big points represent the misclassified samples by the tools. The **A** plot represents the results based on control group, while the **B** plot shows the results based on FBGAN experimental group.

**Fig. S5.**
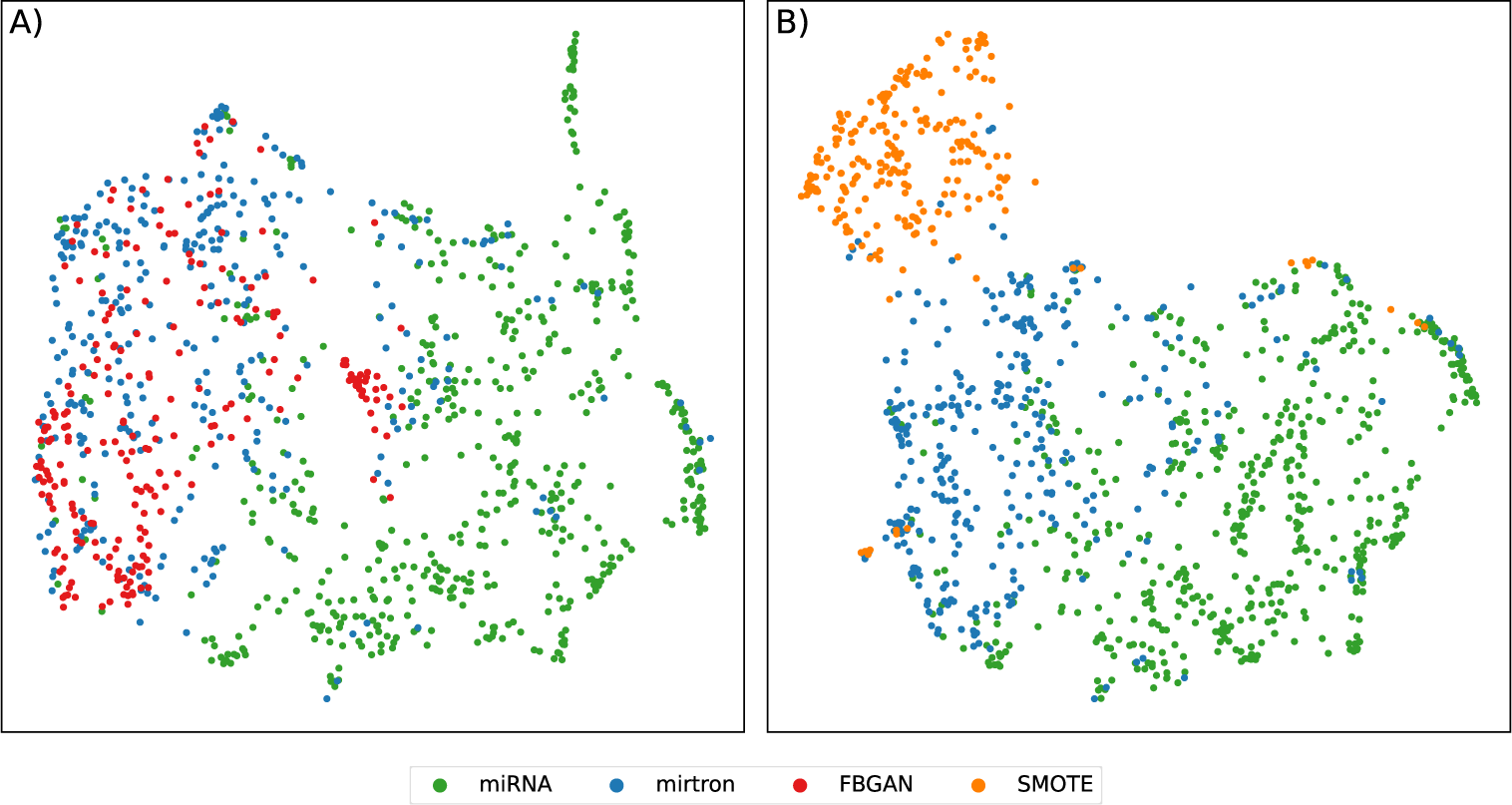
PaCMAP analysis of training sets for FBGAN and SMOTEA experimental groups. The **A** plot shows the distribution of the FBGAN experimental group, while the **B** plot shows the SMOTEA group. The miRNA and mirtron categories are represented by green and blue points, respectively, and the synthetic data points are represented in red (**A**) and orange (**B**).

**Fig. S6.**
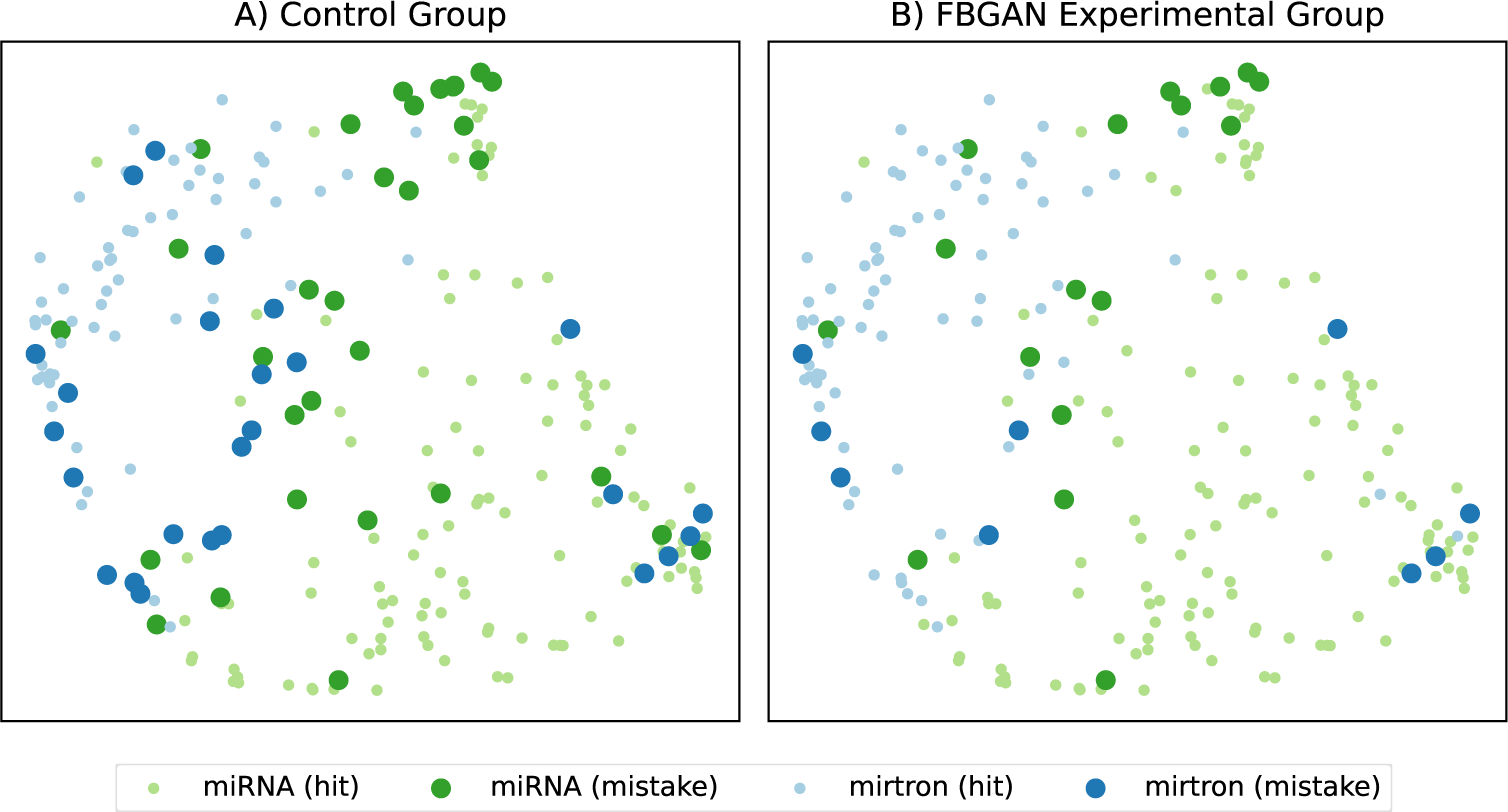
PaCMAP analysis of test set and the classification results of all tools. Both the plots show the two-dimensional space of test set calculated by PaCMAP. The green and blue points represent the miRNA and mirtrons data, and the big points represent the misclassified samples by the tools. The **A** plot represents the results based on control group, while the **B** plot shows the results based on FBGAN experimental group.

### 10.5 Classification performance on other species (Experiment IV)

**Fig. S7.**
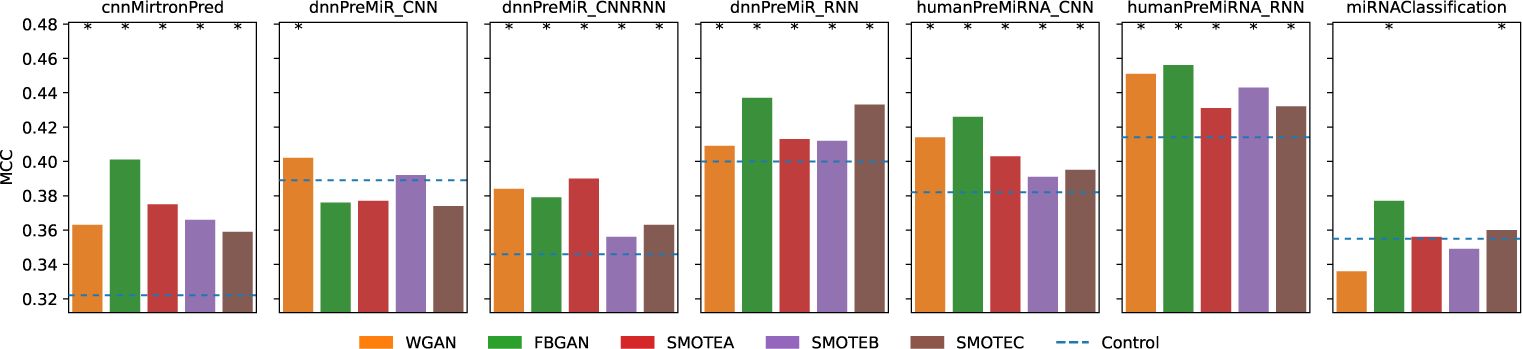
Results for Experiment IV using Dataset II (*H. sapiens*). Each subplot corresponds to a different classification tool, and the colored bars represent the MCC of the classification by the corresponding model of the experimental groups. The blue horizontal line indicates the MCC of the control group for each classification tool. The values above the bars show the improvement in MCC compared to the control. An asterisk (*) above a bar denotes an improvement ≥ 1%.

For the Dataset II (*H. sapiens*), 12 out of 14 (≈ 85.7%) models trained with GAN data showed superior results compared to the control group model (Figure S7). Of the models trained with SMOTE, 16 out of 21 (≈ 76.2%) showed improvement. Only the dnnPreMiR CNN and miRNAClassification tools had results below the control group. When trained with synthetic data, the cnnMirtronPred models achieved the highest MCC increase.

**Fig. S8.**
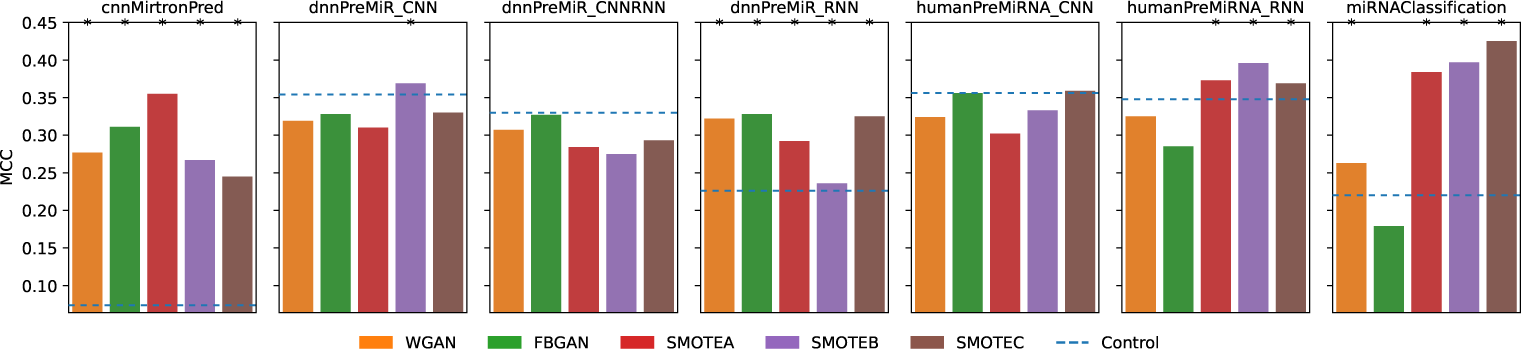
Results for Experiment IV using Dataset III (*M. mulatta*). Each subplot corresponds to a different classification tool, and the colored bars represent the MCC of the classification by the corresponding model of the experimental groups. The blue horizontal line indicates the MCC of the control group for each classification tool. The values above the bars show the improvement in MCC compared to the control. An asterisk (*) above a bar denotes an improvement ≥ 1%.

The classification of Dataset III (*Macaca mulatta* samples) (Figure S8) by the cnnMirtronPred and miRNAClassification tools showed distinct results compared to the other tools. Models of these two tools showed an improvement in generalization ability (except for the miRNAClassification model trained with FBGAN synthetic data). In contrast, the other tools showed little or no improvement in generalization when using synthetic data for model training. Once again, cnnMirtronPred had the highest increase in generalization ability.

#### 10.6 GAN Components

**Fig. S9.**
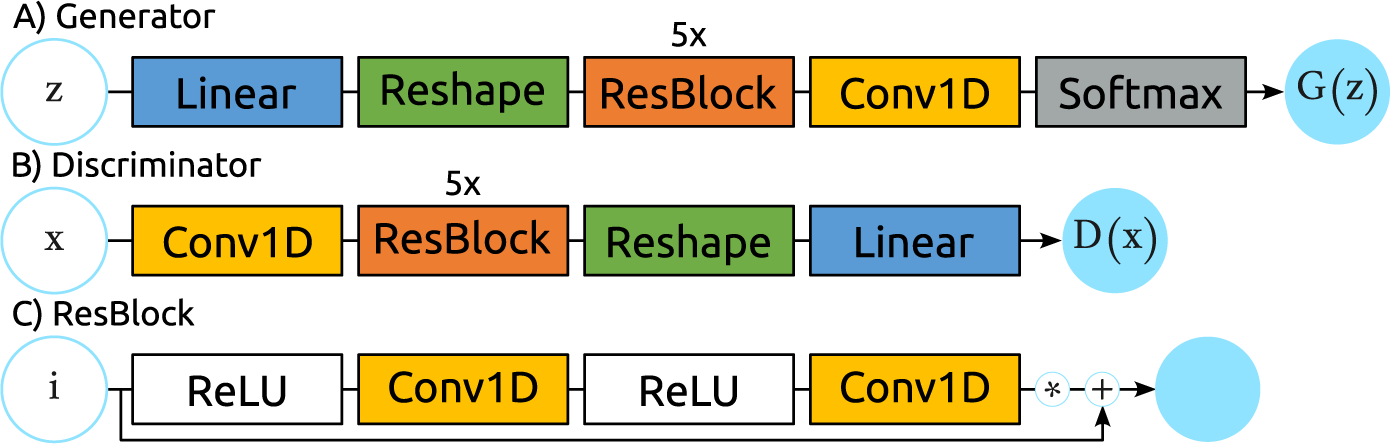
Components of WGAN-GP and FBGAN of this work. **(A)** Generator; **(B)** Discriminator and **(C)** ResBlock used by Discriminator and Generator.

**Table S7.** Discriminator architecture.

|  | Layer | Output shape | Output shape (applied values) |
| --- | --- | --- | --- |
|  | Input | [batch_size, seq_len, n_chars] | [32, 121, 5] |
| 1 | Transpose | [batch_size, n_chars, seq_len] | [32, 5, 121] |
| 2 | Conv1D<sup>1</sup> | [batch_size, hidden_layers, seq_len] | [32, 512, 121] |
| 3 | 5×ResBlock<sup>2</sup> | [batch_size, hidden_layers, seq_len] | [32, 512, 121] |
| 4 | Reshape | [batch_size, hidden_layers * seq_len] | [32, 61952] |
| 5 | Linear | [batch_size, 1] | [32, 1] |

<sup>1</sup> in=n\_chars, out=hidden\_layers, kernel=1
<sup>2</sup> dim\_size=hidden\_layers

**Table S8.** Generator architecture.

|  | Layer | Output shape | Output shape (applied values) |
| --- | --- | --- | --- |
|  | Input | [batch_size, 100] | [32, 100] |
| 1 | Linear | [batch_size, hidden_layers * seq_len] | [32, 61952] |
| 2 | Reshape | [batch_size, hidden_layers, seq_len] | [32, 512, 121] |
| 3 | 5×ResBlock<sup>1</sup> | [batch_size, hidden_layers, seq_len] | [32, 512, 121] |
| 4 | Conv1D | [batch_size, n_chars, seq_len] | [32, 5, 121] |
| 5 | Transpose | [batch_size, seq_len, n_chars] | [32, 121, 5] |
| 6 | Reshape | [batch_size * seq_len, n_chars] | [3872, 5] |
| 7 | Softmax<sup>2</sup> | [batch_size * seq_len, n_chars] | [3872, 5] |
| 8 | Reshape | [batch_size, seq_len, n_chars] | [32, 121, 5] |

<sup>1</sup> dim\_size=hidden\_layers
<sup>2</sup> gumbel\_softmax( $\tau = 0.5$ )

**Table S9.** ResBlock(dim size) architecture.

|  | Layer | Output shape | Output shape<br>(applied values) |
| --- | --- | --- | --- |
|  | Input | [batch_size, dim_size, seq_len] | [32, 512, 121] |
| 1 | ReLU | [batch_size, dim_size, seq_len] | [32, 512, 121] |
| 2 | Conv1d <sup>1</sup> | [batch_size, dim_size, seq_len] | [32, 512, 121] |
| 3 | ReLU | [batch_size, dim_size, seq_len] | [32, 512, 121] |
| 4 | Conv1d <sup>1</sup> | [batch_size, dim_size, seq_len] | [32, 512, 121] |

<sup>1</sup> in=dim\_size, out=dim\_size, kernel=n.chars, padding=2

**Table S10.**
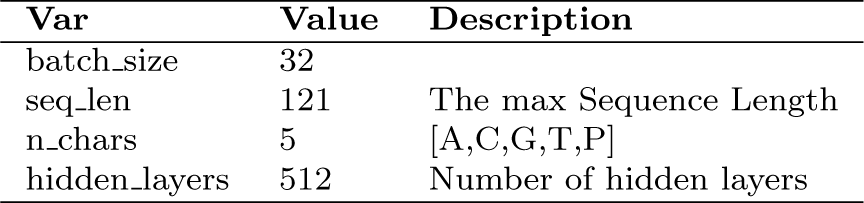
Hyperparameters used in the training of the GANs.

## Notes

### Competing Interest Statement

The authors have declared no competing interest.

